# Laboratory evolution identifies elongated flavodoxins that support electron transfer to sulfite reductases

**DOI:** 10.1101/2023.02.21.529384

**Authors:** Albert Truong, Dru Myerscough, Ian Campbell, Josh Atkinson, Jonathan J. Silberg

## Abstract

Flavodoxins (Flds) mediate the flux of electrons between oxidoreductases in diverse metabolic pathways. While dozens of Fld-partner oxidoreductases have been discovered, these only represent a subset of the oxidoreductases that couple with ferredoxin (Fd) protein electron carriers. To investigate whether Flds can support electron transfer to a sulfite reductase (SIR) that evolved to couple with a Fd, we evaluated the ability of Flds to transfer electrons from a Fd-NADP reductase (FNR) to a Fd-dependent SIR using growth complementation of a microbe with a sulfur metabolism defect. We show that Flds from cyanobacteria complement the growth of this microbe when coexpressed with an FNR and an SIR that evolved to couple with a plant Fd. To better understand the interaction of Fld with these partner oxidoreductases, we evaluated the effect of peptide insertion on Fld-mediated electron transfer. We observe a high insertion sensitivity within regions predicted to be proximal to the cofactor and partner binding sites and a high insertion tolerance within the loop that is used to differentiate short- and long-chain flavodoxins. These results represent the first evidence that Flds can support electron transfer to assimilatory SIRs, and they suggest that the pattern of peptide-insertion tolerance is influenced by interactions with oxidoreductase partners in electron transfer pathways.

## INTRODUCTION

Flavodoxins are low potential electron carriers that use a flavin mononucleotide (FMN) cofactor to transfer electrons between partner oxidoreductases^1^. Biochemical and genetic studies have shown that Flds couple with diverse oxidoreductases possessing cellular roles in glycolysis (pyruvate Fld oxidoreductase, pyruvate formate lyase activating enzyme)^2,3^, photosynthesis (photosystem I)^4^, hydrogen metabolism (hydrogenase)^5^, amino acid synthesis (glutamate synthase, cobalamin-dependent methionine synthase)^6,7^, nucleotide metabolism (ribonucleotide reductase)^8^, steroidogenesis (cytochrome P450)^9,10^, isoprenoid biosynthesis(4-hydroxy-3-methylbut-2-enyl diphosphate synthase) ^11^, redox homeostasis (Fd-NADP reductase, Fld-quinone reductase)^12,13^, lipid synthesis (acyl lipid desaturase)^14^, nitrogen assimilation (nitrogenase, nitrate reductase, nitrite reductase)^6,15–17^, and sulfur metabolism (bisulfite reductase, biotin reductase)^18,19^. In some organisms, Fld deletions result in growth defects, indicating the essential role of Flds in supporting metabolism^20^. In addition, bioinformatic studies have revealed that some organisms have genomes with as many as ten Fld paralogs^21^. Currently, it is not clear why some organisms encode so many Fld paralogs. Some Flds efficiently deliver electrons to many acceptor proteins, including non-natural partners^9,10,22–24^. Other Flds appear to have evolved structures that enable discrimination of partner proteins^10,16,25–28^. Currently, we do not understand how Flds paralogs evolve structures through mutation and elongation to support specialized coupling between different oxidoreductases and pathways.

Many genomes encoding Flds also have ferredoxin (Fd) protein electronic carriers^21^, which function as cellular electron transfer (ET) hubs. The relative abundances of Fld and Fd protein electron carriers can vary widely. Gammaproteobacteria frequently have multiple Fld and Fd paralogs, while microbes from other taxonomic groups typically present higher abundances of Fd electron carriers^21^. Both Fds and Flds present low midpoint reduction potentials. Fld redox potentials range from -230 to -530 mV^29,30^, 2Fe-2S Fds present potentials range from - 150 to -500 mV, and 4Fe-4S Fds range from -200 to -650 mV^31^. In many organisms, Fld and Fd expression is controlled by environmental conditions^32,33^, such as oxidative stress, salt stress, heavy metal toxicity, mineral availability, and light. Also, there is evidence that Fld and Fd expression levels are strongly coupled to iron availability^1,34,35^, with Fld levels being elevated under low iron availability conditions. This trend is thought to arise as iron limitation decreases the availability of substrates required for the biogenesis of iron-sulfur cluster cofactors on Fds^1^. While Fds and Flds support ET to an overlapping set of almost twenty partner proteins, Fds have been shown to couple with a much larger set (>80) of partner oxidoreductases^1,31^. The extent to which Flds can support ET to many of these Fd-partner proteins in natural or synthetic cellular systems remains unclear.

Flds and Fds both support ET to nitrite reductases (NIR)^36^, which share structural similarity with Fd-dependent SIRs^37^. This observation led us to investigate whether Flds could support ET to Fd-type SIRs. Through genome mining, we previously identified microbes that encode Fd, Fd-dependent SIR, and Flds in their genomes^38^. Herein, we show that cyanobacterial Flds support ET between a plant FNR and SIR using a cellular selection that requires ET from a FNR to a Fd-dependent SIR to complement a growth defect^39^. Since some Fld paralogs evolved elongated structures to support recognition of partner proteins^40^, we use this selection to investigate where a cyanobacterial Fld tolerates insertions within its primary structure. We chose this class of mutational lesion because we hypothesized that it would only be tolerated within regions distal from residues that mediate molecular interactions with partner proteins. By comparing peptide-insertion sensitivity profiles with structural models of Fld and Fld-partner complexes, we show that the pattern of Fld peptide-insertion sensitivity correlates with proximity to the cofactor binding site and predicted partner interfaces. Also, we find that the loop that differentiates short- and long-chain Flds presents a high tolerance to peptide insertion^40^. These provide fundamental insight into the ways that Fld evolution could affect oxidoreductase interactions and implicate peptide-insertion profiling as a strategy to rapidly map oxidoreductase binding interfaces. They also identify new Fld-partner proteins that can be used as living electronic components for bioelectronics and synthetic biology^41,42^.

## RESULTS

### Flds support ET to Fd-dependent SIRs

In many microbes, sulfur assimilation from sulfite requires a three-component electron transport chain made up of an electron donor (FNR) that draws reducing equivalents from NADPH, a small protein electron carrier (Fd) that transfers electrons, and a Fd-dependent sulfite reductase that catalyzes the six-electron reduction of sulfite to sulfide^43,44^. To investigate whether Flds might have evolved to substitute for Fds in this pathway, a prior study used bioinformatics to identify organisms encoding Flds as well as this three-component pathway^38^. Photosynthetic cyanobacteria and algae from over ten genera were identified that contain all of these proteins (Fld, Fd, FNR, and SIR), including *Acaryochloris, Anabaena, Crocosphaera, Gloeothece, Nostoc, Ostreococcus, Prochlorococcus, Synechococcus, Thalassiosira*, and *Trichodesmium*. All of these organisms contained a single Fd-dependent SIR, although many contained more than one Fld and Fd. These results suggest that Flds may be capable of supporting electron transfer from FNR to Fd-dependent SIRs.

In prior studies, an *Escherichia coli* strain (EW11) was engineered so that growth is dependent upon Fd-mediated ET from FNR to SIR^39^. This strain cannot grow when sulfate is the sulfur source due to a sulfur metabolism defect, and growth complementation of this defect has been used to report on electron transfer mediated by diverse Fds, including natural pathways made up of corn FNR, Fd, and SIR^39^ as well as synthetic pathways containing non-cognate protein electron carriers^41,45–48^. As such, this strain represents a simple approach to assess whether Flds can support electron transfer between FNR to SIR that evolved to couple with Fds. We first investigated whether *Synechocystis* sp. PCC6803 (sFld1) and *Nostoc* sp. PCC7120 (nFld) Flds, which exhibit 67% identity, could support ET from FNR to SIR (Figure 1a). Each Fld was expressed using an anhydrotetracycline (aTc) inducible promoter (Figure S1a) within a strain also expressing *Zea mays* FNR and SIR using constitutive promoters (Figure S1b) ^45^. To identify optimal assay conditions, we evaluated the growth of cells expressing *Mastigocladus laminosus* Fd and corn FNR and SIR, which complements the growth defect of *E. coli* EW11^45^. When using 96-well plates for this cellular assay, we found that it was critical to use high shaking speeds for robust growth complementation (Figure S1c-e). Under these conditions, Flds complemented cell growth after 48 hours (Figure 1b). In non-selective growth medium, Fld expression had no significant effect on cell growth (Figure S2). These results show that Flds can support cellular ET from an FNR to a Fd-dependent SIR in a synthetic pathway. They also identify growth conditions where Fld ET can be analyzed using a selection.

**Figure 1.**
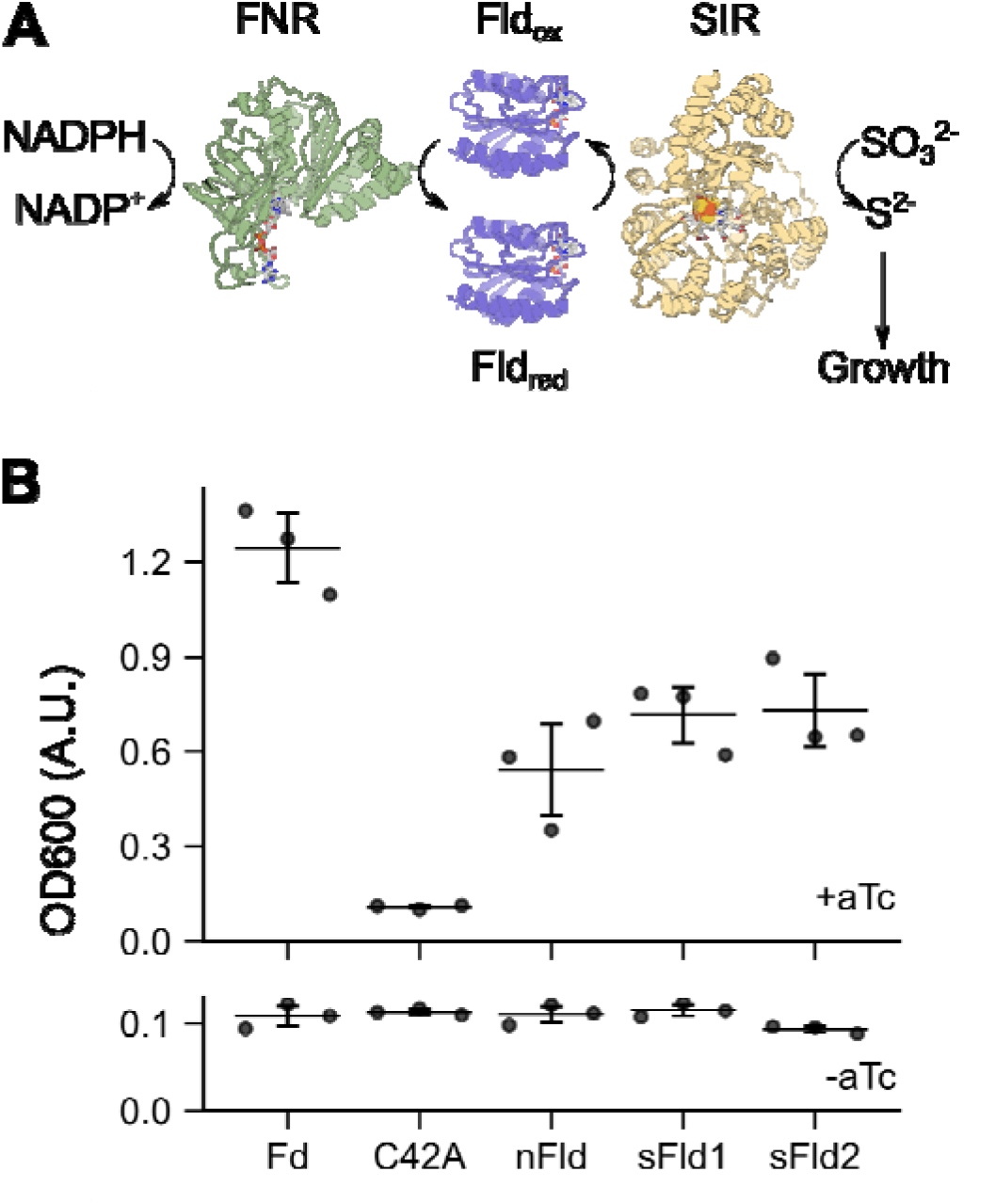
Fld transfer electrons from FNR to SIR. (**A**) The three-component ET pathway tested for growth complementation in *E. coli* EW11. (**B**) *Nostoc* PCC7120 Fld (nFld) and *Synechocystis* PCC6803 Fld (sFld1) both complemented the growth of *E. coli* EW11 after 48 hours at 37°C when the inducer anhydrotetracycline (aTc) was added to growth medium having sulfate as the only sulfur source. As a positive control, we evaluated complementation when cells expressed *Mastigocladus laminosus* (Fd), which has been shown to couple with FNR and SIR. As a negative control, we evaluated complementation using a C42A mutant of this Fd, which cannot coordinate an iron-sulfur cluster. Cells expressing Fd and sFld1 presented significantly higher growth than the negative control (p < 0.02; two-tailed *t* test). While the average nFld OD is ~5-fold higher than C42A, the difference in the mean value of OD from the C42A or uninduced culture only approaches significance (p = 0.0501 and p = 0.0503, respectively; two-tailed *t* test). Cells expressing a Fld from *Synechocystis* PCC6803 containing silent mutations codons 30 and 31 (sFld2) also presented significantly higher growth than this control (p < 0.02, two-tailed *t* test) and do not grow to significantly higher ODs than cells expressing the native sFld1 (p = 0.887). Error bars represent the standard deviation from three biological replicates.

### Evaluating Fld mutation tolerance

Structural studies have shown that some Flds have evolved elongated structures to support high affinity interactions with partner oxidoreductases^21^. While rational design studies have shown that loop removal can be used to disrupt oxidoreductase partner binding^40^, there have been no studies examining how peptide insertion affects Fld electron transfer. To test this idea, we characterized the effects of random peptide insertion on the ability of sFld1 to mediate ET from FNR to SIR. A combinatorial library was built by inserting the octapeptide SGRPGSLS at every backbone location, and the resulting library was selected for Fld-insertion variants that transfer electrons from FNR to SIR using growth complementation of *E. coli* EW11. This peptide was chosen as insertion libraries with this sequence are easy to generate using an existing method^49^. To allow comparison of Fld1 insertion variants with native, we created a plasmid for expressing sFld1 with synonymous mutations, designated sFld2. *E. coli* EW11 expressing sFld1 and sFld2 presented similar growth complementation (Figure 1b). This observation indicates that the barcoded Fld could be embedded in our library selection to calibrate the activities of other Fld mutants.

After building a library of plasmids that express different peptide-insertion variants (Figure S3), we analyzed the library sequence diversity. All of the variants were present in three different sequencing experiments (Figure S4a), with similar average abundances across all variants (Figure 2). For each naive library sequencing experiment (n = 3), the coefficient of variance (CV) for all variants in a sequencing run was calculated. The CVs for each naive library experiment varied from 0.34 to 0.43. We transformed this library into *E. coli* EW11 with sFld2 and analyzed the average sequence diversity before and after selection using three different measurements (Figure S4b-c). Following transformation into this strain, a narrow range of CVs (0.41 to 0.48) was observed for the variant frequencies. The average mutant frequencies in the naive and unselected libraries presented a linear correlation with an R^2^ = 0.82 (Figure S5). With the selected libraries, a larger range of CVs (1.36 to 1.42) was observed within each sequencing run. Taken together, these results show that the sequence diversity in our library was not affected by transformation into *E. coli* EW11, while the selection for growth complementation enriched a subset of variants.

**Figure 2.**
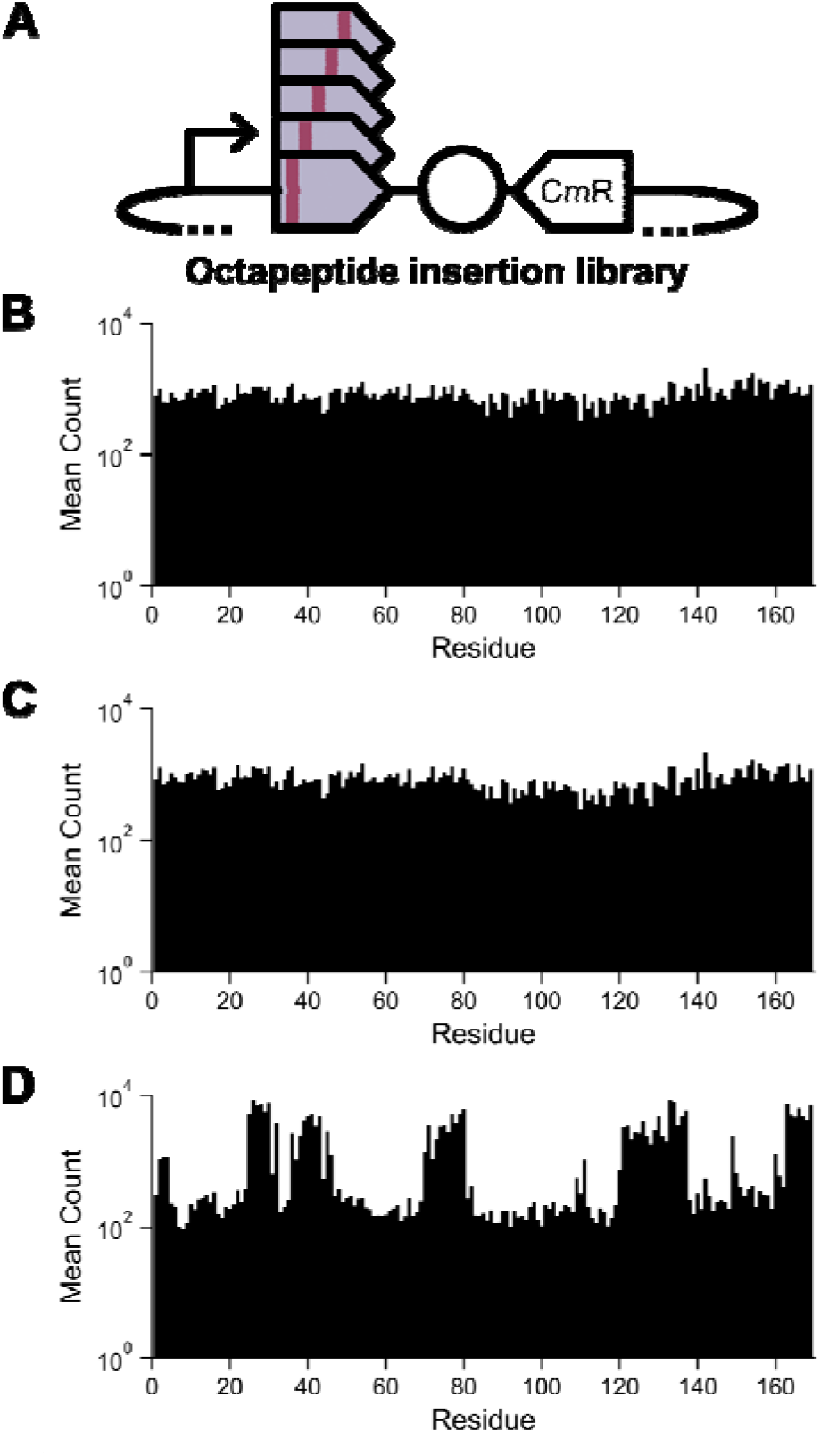
Relative abundances of each peptide-insertion variant. (**A**) A vector library was built to have codons encoding an octapeptide inserted after each codon. Deep sequencing was used to quantify the relative abundances of each Fld variant in the (**B**) naive, (**C**) transformed, and (**D**) selected libraries. For the naive library, the bars represent the mean counts of each insertion variant from three technical replicates, while the selected library represents mean counts from three biological replicates.

To establish which Fld variants support cellular ET, we divided the mean frequency of each variant after selection by the value observed in the naive library (Figure 3a). This analysis yielded *enrichment values* for each variant, defined as the log_2_ (selected:naive ratio). Analysis of the distribution of enrichment values revealed two major clusters of phenotypes, with some variants of intermediate enrichment (Figure 3b). After fitting a three-component gaussian mixture model to the enrichment data, the mean values for the largest modes were -3.00 and 1.45, respectively. The latter cluster overlaps with the enrichment value observed for the sFld2, 1.90 ± 0.14. To determine which variants are non-functional, we evaluated the growth complementation of seventeen peptide-insertion variants (Table S1). As controls, we evaluated cells transformed with a vector that expresses native Fld and a vector that expresses an inactive protein electron carrier. This inactive protein was a Fd-C42A mutant, which does not complement growth^45,50^. After 48 hours (Figure 3c), cells expressing individual peptide-insertion variants presented growth with a linear correlation with fitness scores. These results provide evidence that variants in the peak with low enrichment values are non-functional and that Fld variants with higher fitness scores support cellular ET.

**Figure 3.**
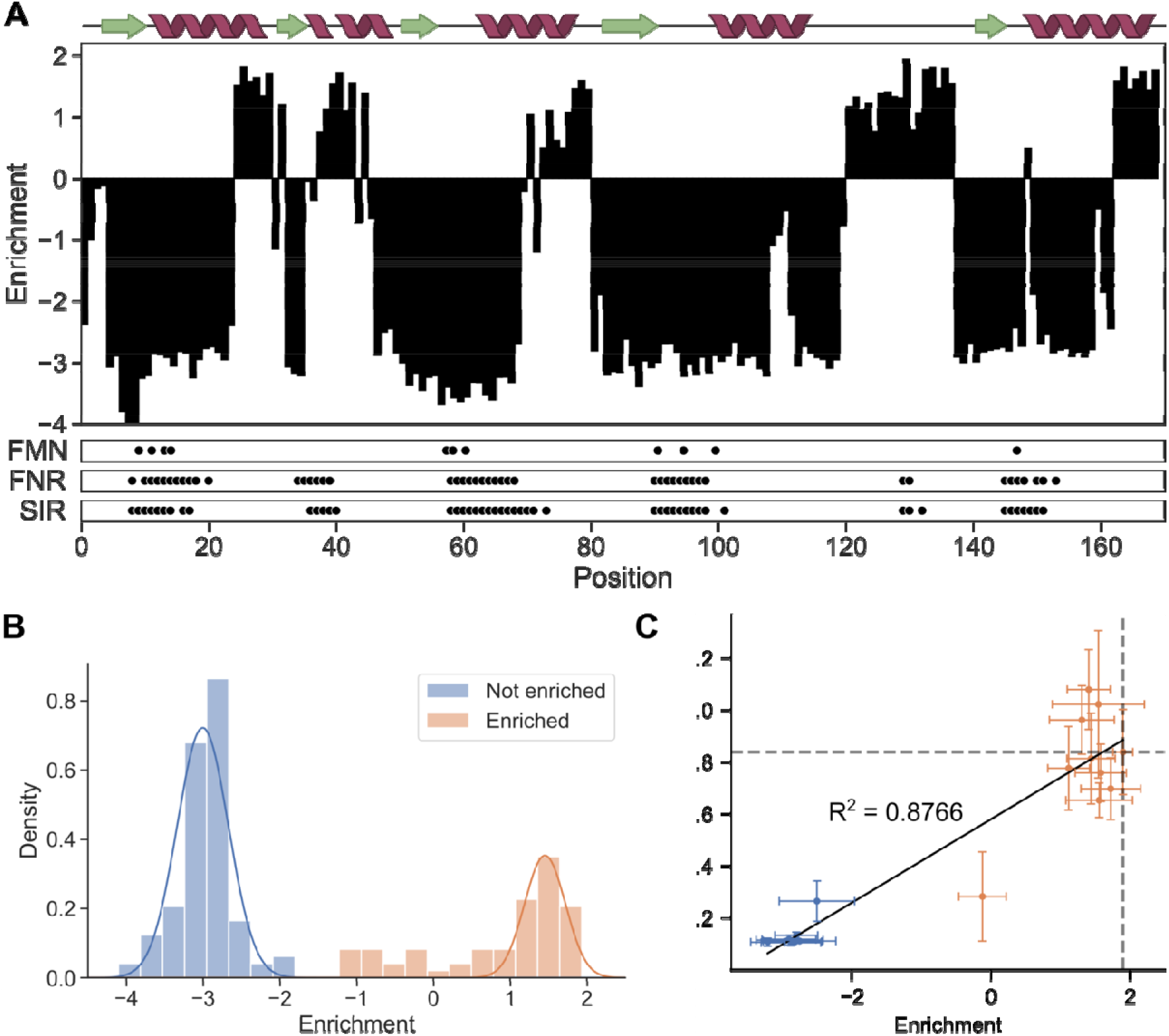
Peptide-insertion profile calculated from sequencing data. (**A**) The relative enrichment of each insertion variant is mapped onto primary structure and compared with predicted secondary structure (top), contacts with the cofactor (FMN), and intermolecular contacts made with the donor (FNR) and acceptor (SIR) partners. Pairs of residues were considered in contact if any constituent heavy atoms were within 8 Å, while enrichment is defined as an enrichment score greater than -1.3. The areas of dots corresponding to interactions between proteins are proportional to the number of residue-residue contacts. (**B**) The distribution of enrichment values fitted to a Gaussian mixture model. Mutants with enrichments within five standard deviations of the low enrichment peak (blue) were designated depleted, while all others having greater values were designated as enriched. (**C**) Comparison of growth complementation of individual Fld variants after 48 hours with calculated enrichment values. A linear fit yields an r^2^ = 0.865. Error bars represent standard deviation from three biological replicates. Dashed lines indicate the mean calculated enrichment value and growth complementation of the sFld2-expressing positive control.

We next created a Fld fitness profile where the enrichment scores were scaled from zero to one, where a value of one represents native Fld activity and a value of zero represents variants with fitness that cannot be distinguished from an empty vector (Figure 4). This profile shows which variants support ET from FNR to SIR. In total, five different Fld motifs tolerated peptide insertion without disrupting Fld-mediated ET from FNR to SIR, including residues spanning from 25 to 30, 39 to 45, 71 to 80, 121 to 137, and 163 to 169. The motif spanning residues 121 to 137 represents the loop that is used to differentiate short- and long-chain Flds^40^. While most other insertion sites presented a high sensitivity to peptide insertion, a handful of sites had intermediate fitness values, including those with peptide insertion after residues 3 to 4, 31, 36 to 37, 44, 46, 72, 109-111, 120, 149, and 160. These findings show that peptide-insertion sensitivity varies with location in Fld primary structure.

**Figure 4.**
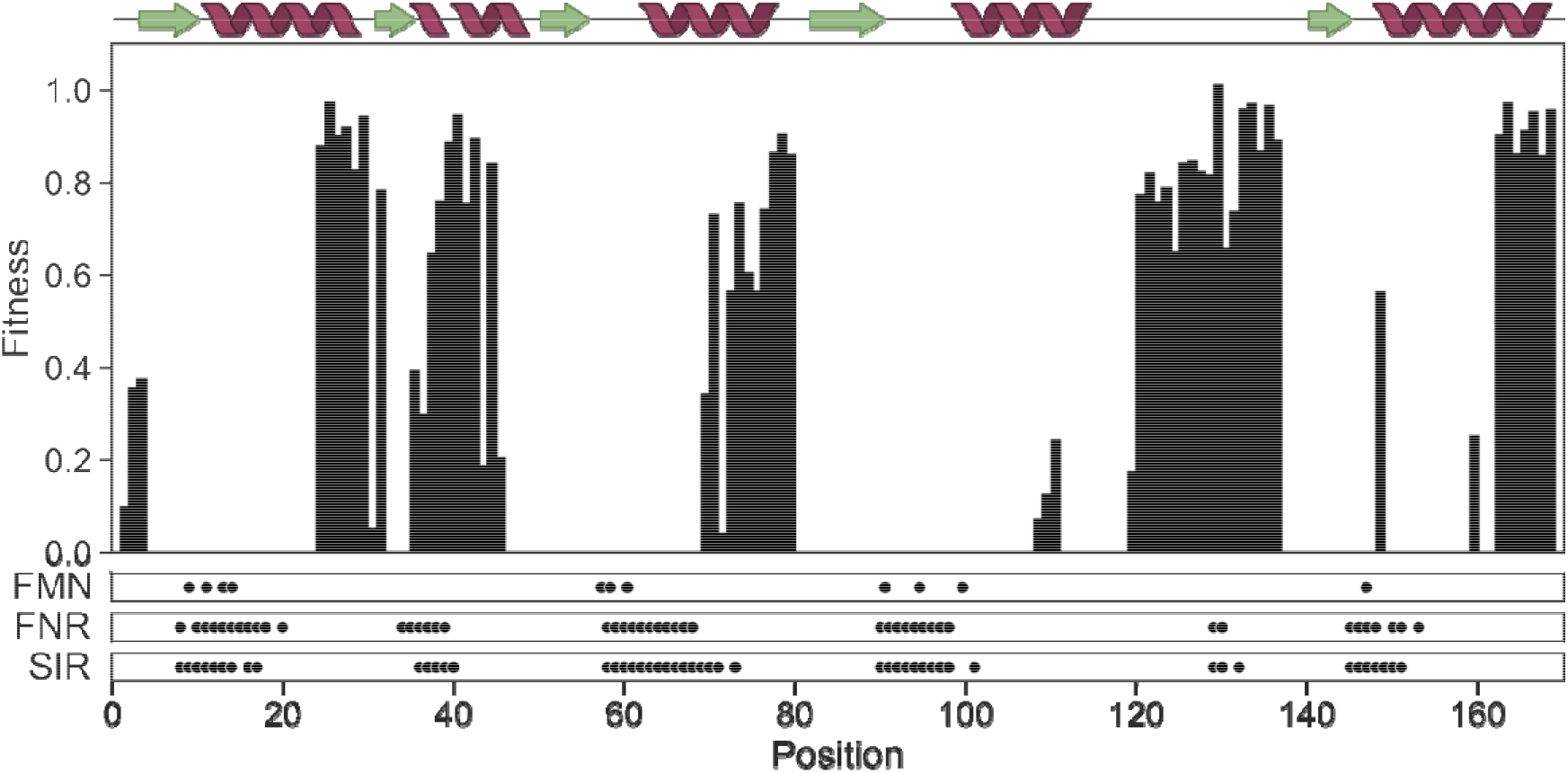
Insertion variant fitness comparison. The fitness of each insertion variant relative to wild type-like Fld is mapped onto primary structure, contacts with the cofactor (FMN), and intermolecular contacts made with the donor (FNR) and acceptor (SIR) oxidoreductase partners.

### Mapping insertion sensitivity onto Fld structure

To better understand the Fld features that influence insertion sensitivity, we compared our results with a sFld1 structural model from AlphaFoldDB^51^. Structural alignment of this Fld with a homolog having 72% sequence identity, *Synechococcus elongatus* Fld^52^, yielded an RMSD of 0.27 Å and revealed similar backbone and side chain conformations of residues at the conserved FMN-binding site (Figure S6). This finding indicated that the FMN binding interface of sFld can be inferred from the crystal structure. When we evaluated how Fld structural features relate to insertion sensitivity, we found that high-fitness variants had insertions in backbone locations that are distal from the FMN (Figure 5A). All peptide-insertion sites proximal to the FMN cofactor (≤ 6Å) were sensitive to peptide insertion (Figure 5B). These findings show that Fld is most sensitive to insertions near the FMN cofactor.

**Figure 5.**
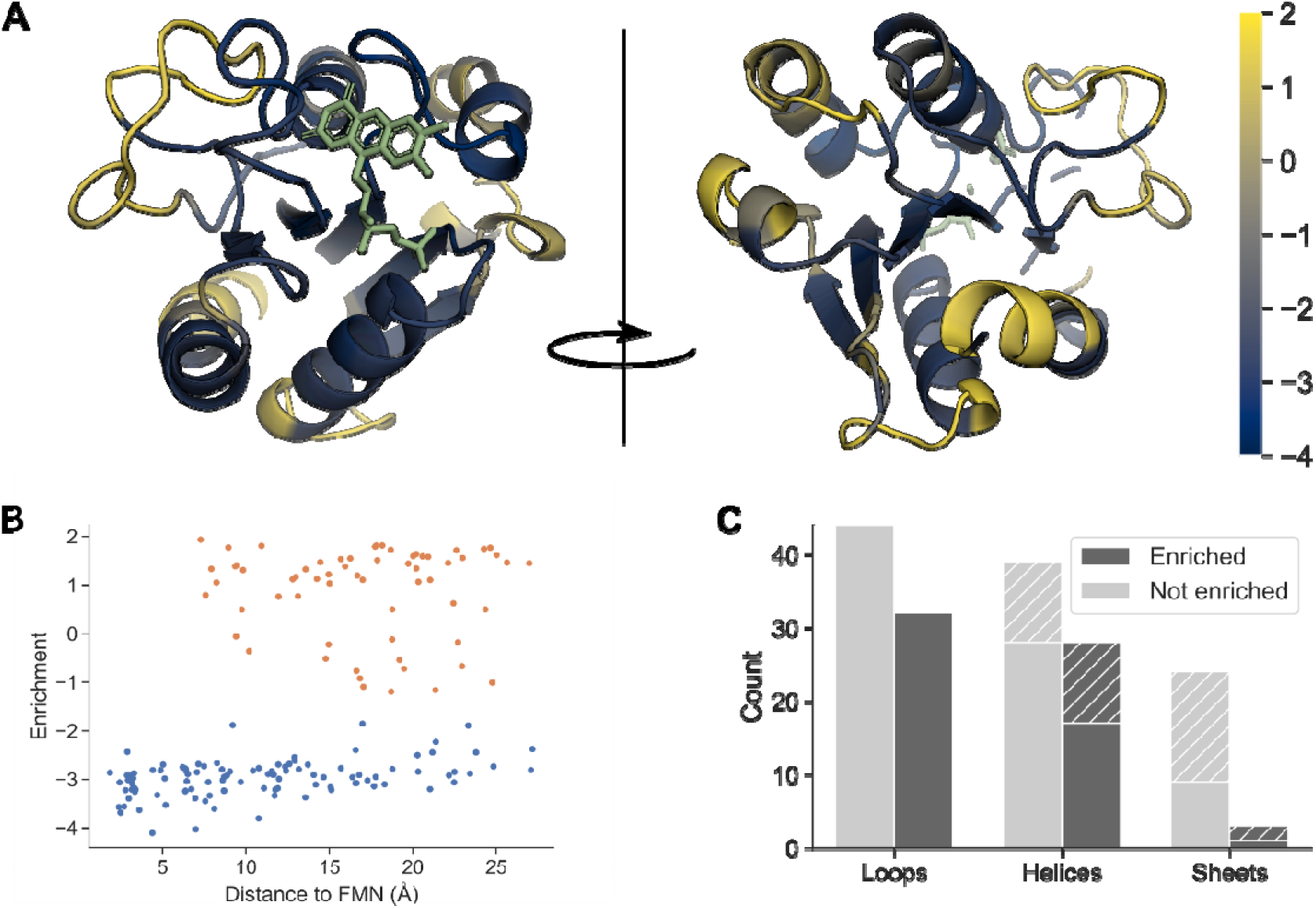
Insertion tolerance correlates with loop proximity and cofactor distance. (**A**) The AlphaFold-predicted structure of the *Synechocystis* Fld colored by enrichment values from the selection. The FMN cofactor was placed by backbone alignment with the crystal structure 1czl. (**B**) Relative enrichments of all variants plotted against the minimum distance between any atom in the residue preceding the insertion site and any atom in the FMN cofactor. (**C**) Relative abundances of enriched and depleted variants by secondary structure. Diagonal lines denote variants having a peptide inserted within two residues of a loop.

We next investigated how insertion sensitivity relates to secondary and tertiary structure. We calculated the number of insertion variants within each secondary structure that support ET (fitness > 0) versus do not support ET (fitness = 0). We observed differential insertion tolerance across secondary structure classes (Figure 5C), with the highest proportion of functional variants resulting from peptide insertion in loops, and the lowest in beta sheets, which are largely found within the core of the protein. We found that many enriched variants with insertions in helices were near loops, so positions within two residues of a loop were considered separately. Of the functional variants within helical and sheet structures, almost half had insertions proximal to a loop. We next evaluated the relationships between fitness and structural features proximal to the insertion site, including contact densities (Figure 6A-B) and residue depth (Figure 6C). We also compared fitness with crystallographic B factors (Figure 6D) from the structure of a homologous Fld to assess whether insertion tolerance may be related to local flexibility. These findings show that native positions that tolerate peptide insertion tend to be less buried and have fewer residue-residue contacts, as well as higher B factors.

**Figure 6.**
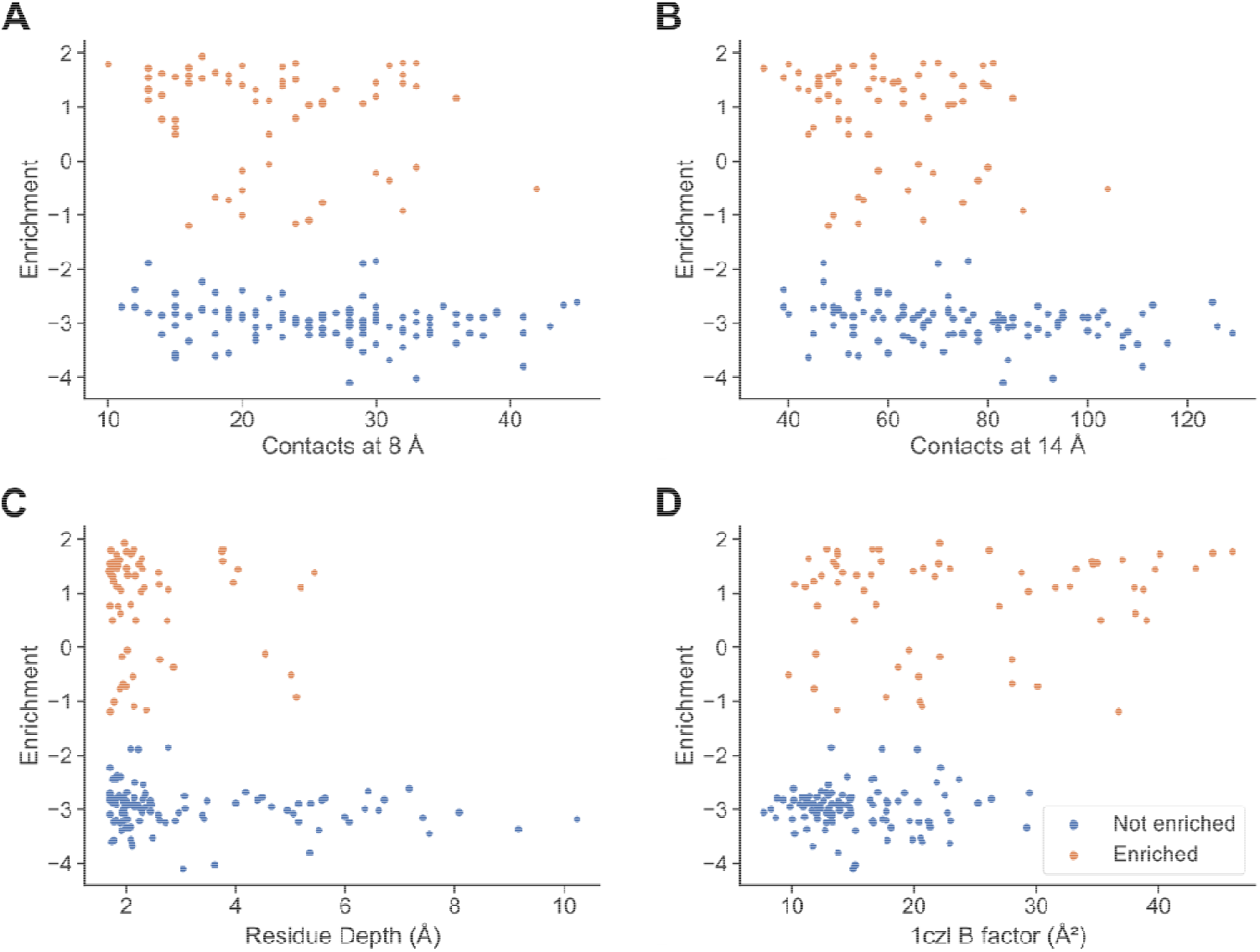
Residues with more intramolecular contacts or greater burial generally have lower insertion tolerance. Relationship between insertion tolerance and residue-residue contact densities at (**A**) 8 Å and (**B**) 14 Å, as well as (**C**) residue depth and (**D**) B factors from the crystal structure of *Synechococcus elongatus* Fld (PDB ID 1czl).

To analyze how peptide-insertion sensitivity relates to partner oxidoreductase binding, we modeled the Fld-FNR and Fld-SIR complexes using AlphaFold-multimer^53,54^. We observed two binding modes for the Fld-FNR complex (Figure S7) and aligned Fld (PDB ID 1czl) and *Z. mays* FNR (PDB ID 1jb9) crystal structures to each prediction to obtain the orientations of the FMN and FAD cofactors. To determine if the predicted conformations are compatible with fast intermolecular ET, we compared the cofactor orientations to those of cytochrome P450 reductases (CPRs) (Figure S8), a protein family comprising fused Fld and FNR domains^55^. One of the predicted Fld-FNR binding modes recapitulated the side-to-side orientation of the FMN and FAD cofactors observed in CPRs with an interatomic distance of 3.2 Å, which is conducive to intermolecular ET. When we analyzed how peptide-insertion sensitivity correlates with Fld-FNR intermolecular contact densities in the modeled structural complex (Figure 7A), we found that sensitivity was related to intermolecular contact density (Figure 7B). All Fld variants with insertions at positions making ≥ 5 contacts with FNR were non-functional. In regions making 0 to 4, the insertion tolerance was bimodal. This finding suggests that the pattern of peptide-insertion sensitivity contains information about the Fld regions that mediate FNR binding.

**Figure 7.**
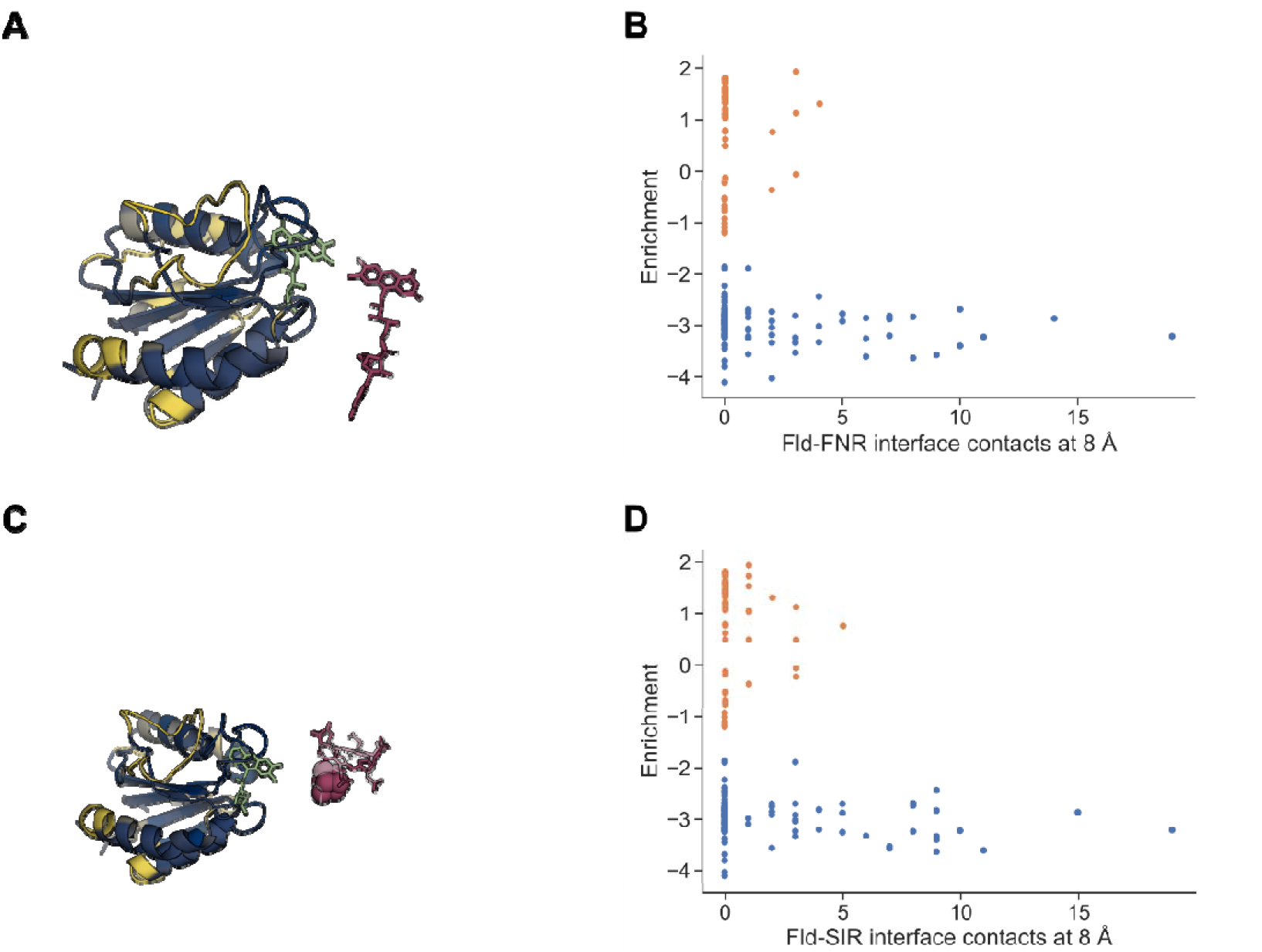
Positions at the predicted interfaces with partner oxidoreductases do not tolerate insertions. (**A**) One of two predicted binding modes predicted for the Fld-FNR complex. This binding mode recapitulates the interflavin geometry observed among cytochrome reductases. (**B**) Enrichments of Fld insertion variants plotted against numbers of intermolecular residue-residue contacts with FNR at 8 Å. (**C**) The predicted binding modes for the Fld-SIR complex. (**D**) Enrichments of Fld insertion variants plotted against numbers of intermolecular residue-residue contacts with SIR at 8 Å. For both complexes, contacts were counted for the residue preceding the insertion.

A Fld-SIR structural model was also created using AlphaFold-multimer. To assess whether the predicted complex is compatible with efficient intercofactor ET, we aligned Fld (PDB ID 1czl) and *Z. mays* SIR (PDB ID 5h92) crystal structures with our Fld-SIR model (Figure S9). The intercofactor distance for the Fld-SIR complex was 8.3 Å, which is slightly lower than the intercofactor distances (~12 Å) observed in crystal structures of the SIR with its native Fd partner^43^. A comparison of the intercofactor distances and orientations in the Fld-SIR and Fd-SIR complexes is shown in Figure S10. We next analyzed how peptide-insertion sensitivity correlates with Fld-SIR intermolecular contact densities in the predicted complex (Figure 7C). As observed with the Fld-FNR complex, peptide-insertion sensitivity correlated with intermolecular contact density in the Fld-SIR complex (Figure 7D). All variants having insertions at Fld locations making ≥ 5 contacts with FNR were non-functional. In regions making 0 to 4 contacts, the insertion tolerance was bimodal. This finding suggests that the pattern of insertion sensitivity contains information about the Fld regions that mediate binding to SIR.

## DISCUSSION

Our results herein extend the list of Fld-partner oxidoreductases to include Fd-dependent SIRs. They also show the utility of synthetic electron transfer pathways for rapidly assessing coupling of protein electron carriers with different partner oxidoreductases^31^. The finding that cyanobacterial Flds can mediate ET from plant FNR to SIR implicate a role for this three-component ET pathway in microbes whose genomes encode all three of these oxidoreductases. Prior bioinformatic analysis revealed that a majority of the microbes containing FNR, Flds and Fd-dependent SIR are marine cyanobacteria, such as *Prochlorococcus* and *Synechocystis*^38^. Biogeochemical studies have shown that iron is a limiting resource for primary productivity in marine settings^56^, and they have shown that cyanobacteria compete for iron with heterotrophic bacteria and phytoplankton^57^. Taken together with the finding that sulfate is abundant in marine settings, 28 mM^58^, our results suggest that Fld-mediated electron transfer from FNR to SIR may occur under conditions when iron is sufficiently limiting so that it cannot support iron-sulfur cluster biogenesis critical to Fd electron transfer^59^. To directly demonstrate a physiological role for the Fld-SIR interactions, studies will be needed to directly show coupling with cognate FNR and SIR under physiological conditions.

The finding that insertion sensitivity depends on proximity to cofactor and partner binding interfaces illustrates how peptide-insertion tolerance could be used to map residues that mediate molecular interactions in the absence of structural information. Peptide-insertion sensitivity correlates strongly with three structural parameters, including the density of intramolecular contacts, proximity to cofactor, and the density of intermolecular contacts. Prior studies have probed protein structure by inserting peptides with a range of lengths^60–64^. These studies found that peptide insertion can provide insight into molecular interactions^60^, conditional phenotypes for genetic studies^61^, and sites compatible with the creation of non-disruptive affinity tags^62^. A major limitation of these prior studies is the use of transposon mutagenesis to generate sequence diversity, which results in non-uniform variant abundances within naive libraries^65^. Our study leveraged a recently described algorithm for domain insertion to create more uniform libraries^66^, which enabled us to create a comprehensive fitness map. By scoring the effects of peptide insertion on cellular function, we observed strong correlations between insertion sensitivity and intramolecular and intermolecular interactions. Further studies will be required to establish the mechanisms by which different sequence changes affect function, *e*.*g*., direct measurements of folding, cofactor binding, electrochemical midpoint potential, and partner-binding.

Insertions and deletions contribute to protein evolution, but it remains unclear how structural and functional features constrain changes in sequence length^67^. Similar to a prior study that examined protein tolerance to single amino acid insertions^68^, we found that the insertional tolerance correlates strongly with the identity of the secondary structure targeted by insertion, with loop tolerance being greater than that of helices and helices having a greater tolerance than beta strands, and we observed a correlation with insertion tolerance and proximity to the end of helix and strand secondary structures. Our study extends these findings by showing a correlation between insertion tolerance and proximity to cofactor and partner protein binding sites. In a recent study, peptide-insertion sensitivity also correlated with cofactor proximity within a membrane protein wire^64^. Taken together, these results support the idea that tolerance to insertion is constrained by the protein fold, requiring considerations of nonlocal interactions. Moreover, insertion tolerance at a given site varies based on the size and characteristics of the inserted peptide^69^, which may display its own secondary and tertiary structure preferences. At times, these perturbations can lead to alternate frame folding^70^. Such dramatic changes in protein structure create challenges for predicting the effects of insertions from native structures computationally except in specific cases^71^. Deep learning approaches offer promise for predicting fitness directly from sequence^72–74^. However, they require larger insertion datasets for refinement and validation and to reveal principles governing insertion tolerance^75^. This can be contrasted with deep mutational scanning experiments that study the effects of amino acid substitutions^76^. The fitness effects of random mutations provide insight into how individual residues contribute to folding, stability, dynamics, and function. Amino acid preference at a given site in a protein is largely determined by the local molecular environment, as reflected by the recent success of deep learning approaches for variant prediction and sequence recovery from structure^77–79^, rather than non-local interactions of importance for insertions.

Our results illustrate how insertional mutagenesis can be used to understand how proteins have evolved elongated structures, which has occurred through the recombination of protein domains^80^. It is unclear how the identity of the protein homolog targeted for random peptide insertion will affect insertion tolerance, as well as the partner oxidoreductases used to select for coupling. Prior studies varying the stability of the protein homolog targeted for mutagenesis have revealed that sensitivity to mutations decreases as the targeted protein increases in thermostability^81–83^. Peptide insertion profiles created with multiple homologs will be useful for establishing how topology, stability, and local energetics govern mutational tolerance^84^. Also, it will be interesting to use this approach with peptides of varying sequences and lengths to generate larger data sets as a means to develop rules for domain insertion^85^. Emerging machine learning approaches can be applied to these data sets in tandem with metagenomic data to predict design rules for elongated proteins^69,74^. Such rules will be critical for rationally inserting larger polypeptides as a means to create protein switches for synthetic biology^86^, such as polypeptides encoding ligand-binding domains. The creation of such allosteric switches within protein electron carriers is needed to dynamically regulate electron flow in living sensors created for metabolic flux control and sensing applications^41,45^.

## METHODS

### Materials

Chemicals were from VWR, MilliporeSigma, Fisher, Apex Biosciences, Research Products International, or BD Biosciences. Enzymes and molecular biology kits were from Zymo Research, Qiagen, and New England Biolabs.

### Strains

*E. coli* XL1-Blue (Agilent, Inc) and Turbo Competent *E. coli* (New England Biolabs) were used for plasmid construction, E. cloni 10G (Lucigen) was used for library construction, and *E. coli* EW11 was used for growth complementation^39^. For molecular biology, cells were grown at 37 °C while shaking at 250 rpm in lysogeny broth (LB) pH 7. Cells were transformed using electroporation (1 pulse, ~5 ms, 1.8 kV, 1 mm gap electroporation cuvette). Following transformation, cells were allowed to recover in super optimal broth (SOB) pH 7 for 1 hour at 37 °C, which contained 5 g/L yeast extract, 20 g/L tryptone, 10 mM sodium chloride, 2 mM potassium chloride, and 20 mM magnesium sulfate.

### Plasmids

Table S2 lists the plasmids used in this study. *Zea mays* FNR and SIR were constitutively expressed using pSAC01^45^. The control vectors expressing *Mastigocladus laminosus* Fd (pFd007) and an inactive C42A mutant of this Fd^45,50^. Plasmids for expressing sFld1 and nFld under control of aTc-inducible promoter (pAG034 and pAG036, respectively) were created by PCR amplifying commercially synthesized genes (Integrated DNA Technologies, Inc.) and cloning them into pFd007^45^, which contains a ColE1 origin, chloramphenicol resistance gene, and a synthetic translation initiation region. A plasmid for expressing sFld2, which encoded sFld1 with a barcode, was created with synonymous mutations in the codons for residues 30 and 31 (designated pAT001); these codons were mutated from AGTGTG to TCCGTT. To create plasmids for assessing the growth complementation of individual peptide-insertion mutants, pAG034 was PCR amplified using primers that code for insertion and circularized using Golden Gate^87^. Mutant plasmids were also isolated from the naive peptide-insertion library. All plasmids were sequence verified.

### Growth complementation

ET from FNR to SIR was evaluated using growth complementation of *E. coli* EW11 as previously described^39^. To perform growth complementation, cells were transformed with pSAC01^45^ and plasmids expressing protein electron carriers. Individual colonies were grown in a modified m9 minimal medium (m9c) that includes both sulfur-containing amino acids (80 mg/L each), as well as sodium phosphate heptahydrate, dibasic (6.8 g/L), potassium phosphate, monobasic (3 g/L), sodium chloride (0.5 g/L), 2% glucose, ammonium chloride (1 g/L), calcium chloride (11 mg/L), magnesium sulfate (0.24 g/L), ferric citrate (0.12 g/L), p-aminobenzoic acid (2 mg/L), inositol (20 mg/L), adenine (5 mg/L), uracil (20 mg/L), tryptophan (40 mg/L), tyrosine (1.2 mg/L), and the remaining 16 amino acids (80 mg/L each). To evaluate complementation, cells were transferred to a m9 minimal medium that is selective (m9sa). This medium is identical to m9c other than the lack of sulfur-containing amino acids, with magnesium sulfate (0.24 mg/L) as the only S source.

### Library synthesis

The peptide-insertion library was created as described previously^49^. In brief, the sFld1 gene was computationally fragmented into four tiles, and variants of each tile were created that contained an 8-codon insertion between every codon in the tile. Tile sequences, as well as primer sequences for cloning, were generated and synthesized by Twist Biosciences and Integrated DNA Technologies, respectively. Each tile was PCR amplified using Q5 DNA polymerase, the amplicons were gel purified, and Golden Gate cloning with BsmBI was used to insert each tile into pAG034, a vector with a chloramphenicol resistance marker that expresses sFld1 under the control of an aTc-inducible promoter. Each vector ensemble was transformed into E. cloni 10G and plated on LB-agar medium containing chloramphenicol (34 ng/mL). After overnight incubation at 37 °C, colony forming units (CFU) were quantified. Each vector ensemble, which contained ~50 variants, yielded ≥ 3600 CFU. Assuming that ~50% of the synthesized oligo library sequences contain errors and sampling replacement, this colony count indicates that >99% of our variants were sampled at this step^88^. To generate the naive library, colonies were harvested from plates, pooled, and purified using a miniprep kit (Qiagen, Inc.). Library sequence diversity was evaluated using AmpliconEZ sequencing (Genewiz, Inc.).

### Library Selection

Electrocompetent *E. coli* EW11 containing pSAC01 were transformed by electroporation with a mixture of the naive library (22.3 fmols) and the barcoded sFld2 vector pAT001 (1.2 fmols). Following transformation, cells were grown in SOB while shaking at 250 rpm for 1 hour at 37 °C. The culture was split into three equal volumes, which were plated on LB-agar medium containing chloramphenicol and streptomycin, 34 ng/mL and 100 ng/mL, respectively. Plates were incubated overnight at 37 °C, CFU were visually counted (29,500 CFU). To prepare glycerol stocks of resulting transformants for this unselected library, plates were scraped, and a cell pellet was obtained by centrifugation for 5 minutes at 4000g. The cells were washed twice by resuspending the pellet in 50% glycerol followed by centrifugation. Washed cells were resuspended in 50% glycerol, divided into aliquots, and stored at -80 °C. Prior to use, the CFU/mL of the cell library was calculated by plating serial dilutions of library stock on LB-agar plates containing 34 ng/mL chloramphenicol and 100 ng/mL streptomycin. To select the library for Flds that mediate ET from FNR to SIR, *E. coli* EW11 transformed with the library (5 × 10^6^ CFUs) was used to inoculate three flasks containing m9sa medium (30 mL) containing with 34 ng/mL chloramphenicol, 100 ng/mL streptomycin, and 100 ng/mL anhydrotetracycline. The number of CFUs used to inoculate the media was calculated by counting colonies on LB-agar medium containing antibiotics following serial dilutions of the inoculum. The m9sa cultures were then incubated at 37°C while shaking at 300 rpm. After 48 hours, the selected cultures grew to 3.12 × 10^8^, 9.8 × 10^8^, and 3.6 × 10^8^ CFU/mL. Plasmids were isolated from the inoculum and the m9sa cultures following selection using a miniprep kit (Qiagen, Inc.).

### Sequencing

Prior to deep sequencing, each half of the Fld gene in the naive, transformed, and selected libraries was PCR amplified using Q5 DNA polymerase. This PCR amplification added unique sequencing adaptors to the ends of each amplicon (Table S3). To sequence the first half of the Fld gene, adaptors were used that add 42 base pairs prior to the start codon and after base pair 297 in the coding sequence. To sequence the second half, adaptors were added prior to base pair 188 of the coding sequence and after a location 39 base pairs following the stop codon. All of these amplicons were sequenced using the Genewiz AmpliconEZ service. Individual plasmids were sequence verified using Sanger sequencing (Genewiz, Inc.). Raw sequence reads, frequency values, and statistical analysis are provided as Supplementary Data.

### Peptide-insertion profile

The paired reads representing the forward and reverse sequencing data were merged using the BBmerge.sh script^89^. Sequencing reads that were unable to be merged were not used in calculations of enrichment. In cases where >90% of reads were unmergeable, the full data set was not used for enrichment calculations. To identify sequences containing an inserted peptide, we used a previously described python script (github.com/SavageLab/dipseq) developed for domain insertion analysis^90^. To eliminate sequences that contained out-of-frame insertions and to graph the abundances of each unique in-frame insertion variant, we used a python script (https://github.com/SilbergLabRice/dipseqplotter). The frequency (F) of each insertion variant in each sequencing run was calculated by dividing the number of identical insertion variant reads by the total number of merged sequencing reads. The enrichment of mutants following selection was quantified by calculating the log_2_ (fold change), where the fold change represents the ratio of the selected to the naive frequencies. When mutants had an enrichment score below -1.3, they were designated inactive. This threshold was obtained by fitting a gaussian mixture model to the distribution of log-enrichment scores. All variants <5 standard deviations higher than the mean of the low enrichment peak (mean = -3.0) were designated as non-functional. Using this scoring, fitness scores ranging from 0 to 1 were assigned, where 1 represents the parental Fld enrichment and 0 represents Fld variants that do not present cellular ET.

### Structural modeling

A model of the sFld1 (IsiB) structure from AlphaFoldDB (entry P27319) was used for structural analysis of free sFld1^51^. All residues except the first and last two residues had confidence metrics (pLDDTs) above 90, and the lowest pLDDT was >70. These high values led us to use all modeled positions for further analysis. The sFld1 exhibits 72% identity to *Synechococcus elongatus* IsiB, a structurally characterized homolog, PDB ID 1czl^52^, enabling identification of residues at the conserved FMN binding site. AlphaFold-multimer was used to model the Fld-FNR or Fld-SIR complexes^53,54^. Target sequences were obtained from Uniprot for the sFld1 IsiB (P27319), *Z. mays* chloroplastic SIR (O23813) and *Z. mays* root FNR (Q41736). To mirror the partner-protein sequences used in our cellular assay, the N-terminal ten residues of the FNR were omitted and a glycine residue was added between Met11 and Ser12^39^. Also, the N-terminal chloroplast localization sequence was omitted when modeling SIR. The primary structures used for all modeling are provided as supporting information. To obtain models for the Fld-FNR and Fld-SIR complexes, we used the ColabFold^91^ implementation of AlphaFold-multimer version 2.2.0 with multiple sequence alignments (MSAs) obtained from ColabFoldDB^91^ using the mmSEQs2 webserver^92,93^. According to the default behavior of AlphaFold/ColabFold, sequences from the same organism were paired in the MSA, and unpaired sequences were also included. Templates were included from the PDB70 version 13Mar22^94,95^. Cofactors were placed in each predicted structure by backbone alignment with structures of Fld PDB ID 1czl^52^, SIR PDB ID 5h92^43^, and FNR PDB ID 1jb9^96^.

The confidence metrics for each AlphaFold protein complex prediction are provided in Table S4. For the Fld-FNR complex, the first binding mode had a predicted template modeling (pTM) score of 0.91 and interface pTM (pTM) of 0.86, while the second binding mode had a pTM score of 0.89 and ipTM of 0.82. Predicted aligned error (PAE) and predicted local-distance difference test (pLDDT) plots are included in Figure S10. We chose the second binding mode for our analysis, despite the lower ipTM, since the orientations of the FNR and Fld as well as their cofactors closely recapitulated the productive binding mode of the Fld and FNR domains of cytochrome reductases. This correspondence did not arise from the use of templates, as AlphaFold-multimer cannot use template information for predicting relative orientations of separate proteins^54^. The intercofactor distances and orientations of the Fld and FNR in this binding mode are similar to those for a pair of homologues using conventional docking approaches^97^.

For the Fld-SIR complex, a single binding mode was predicted with a pTM score of 0.93 and interface pTM (ipTM) of 0.86. The PAE values are higher than the Fld-FNR complexes, which did not improve with additional recycling. Given the high MSA depth, this uncertainty is thought to arise from the shallow, smooth, and highly-charged interface of the corn SIR, which is compatible with multiple binding conformations. Ensemble binding has been observed with the corn SIR and its native ferredoxin partner, which binds in at least three conformations that primarily vary by rigid-body rotation along the inter-cofactor axis^43^.

### Structural calculations

To quantify intramolecular contact density, residue-residue interactions were identified using distance cutoffs of 8 Å and 14 Å, which are commonly used to consider medium-range contacts alone and medium-range and long-range contacts together^98,99^. Intermolecular contacts were considered separately to identify residues at the predicted interfaces with FNR and SIR. Residue depth was calculated using the MSMS package as the average depth over all atoms in a residue^100^. Local flexibility was inferred from the Cα B factors from a related Fld crystal structure^52^.

### Statistics

Library selection data is presented as the mean and standard deviation of three biological (selective library) or technical (naive and non-selected library) replicates. P-values were obtained using a two-tailed Welch’s t-test. A three-component gaussian mixture model was fit to the enrichment data using the Scikit-learn library for Python^101^

## SUPPLEMENTAL MATERIAL

Supplemental Figures S1-S10 and Tables S1-S4 are provided as a single PDF file. Also, sequencing counts and frequencies are provided as a single XLSX file.

## ACKNOWLEDGEMENTS

We are grateful for support from the Office of Basic Energy Sciences of the U.S. Department of Energy grant DE-SC0014462 (to JJS), Office of Naval Research grant N00014-20-1-2274 (to JJS), and NSF Postdoctoral Research Fellowships in Biology Program under Grant No. 2010604 (to JTA).

## Notes

### Competing Interest Statement

The authors have declared no competing interest.

