## Supplemental Figures for "Laboratory evolution identifies elongated flavodoxins that support electron transfer to sulfite reductases"

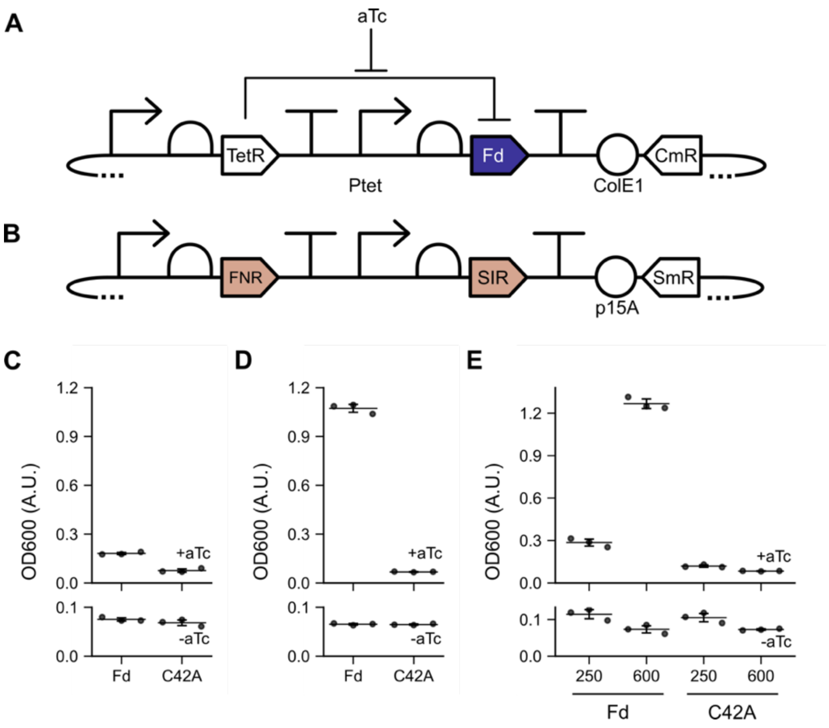


**Figure S1.** Effect of growth conditions on Fd complementation. Vectors for expressing (**A**) Fd and (**B**) corn FNR and SIR. (**C**) Growth complementation in deep 96-well plates containing 1 mL m9sa medium following a 37°C incubation while shaking at 250 rpm for 24 hours. Cells expressing Fd presented an OD that was 2.4-fold higher than cells expressing an inactive mutant (C42A) ($P < 0.01$; two-tailed *t* test). (**D**) Complementation in 14 mL round-bottom Falcon tubes containing 1 mL of m9sa following a 37°C incubation while shaking at 250 rpm for 24 hours. Cells expressing Fd had a ~15.8-fold higher OD than cells expressing C42A ($P < 0.01$; two-tailed *t* test). (**E**) Increasing the shaking speed of 96-well plates from 250 rpm to 600 rpm and decreasing the volume from 1 to 0.5 mL significantly increased the OD of Fd-expressing cells after 24 hours ($p < 0.01$; two-tailed *t* test). Error bars represent $\pm 1\sigma$ from 3 replicates.

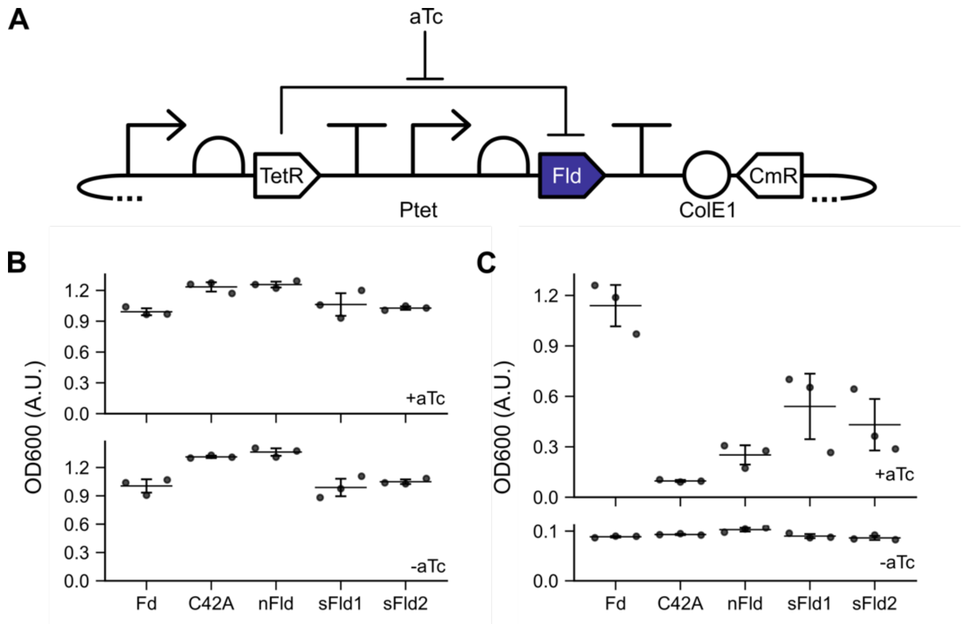


**Figure S2.** Flds present a low fitness burden under non-selective conditions. (**A**) Plasmid used to conditionally express Flds. (**B**) Growth of cells expressing SIR and FNR alongside different protein electron carriers, including a Fd, an inactive Fd mutant (C42A), *Nostoc* PCC7120 Fld (nFld), *Synechocystis* PCC6803 Fld (sFld1), and sFld1 harboring silent mutations at codons 30 and 31 (sFld2). Growth was not affected by addition of an inducer (aTc) for expressing the protein electron carriers following a 20 hour incubation in nonselective m9c medium ($p > 0.05$, two-tailed *t* test), except for cells expressing nFld, in which a small but significant 1.09-fold decrease in OD was observed between uninduced and induced cultures ($p = 0.04$, two-tailed *t* test). (**C**) When an identical experiment was performed in selective medium, only the cells expressing Fd grow to significantly higher OD than cells that express C42A ($p < 0.01$, two-tailed *t* test). Error bars represent $\pm 1\sigma$ from 3 replicates.

.

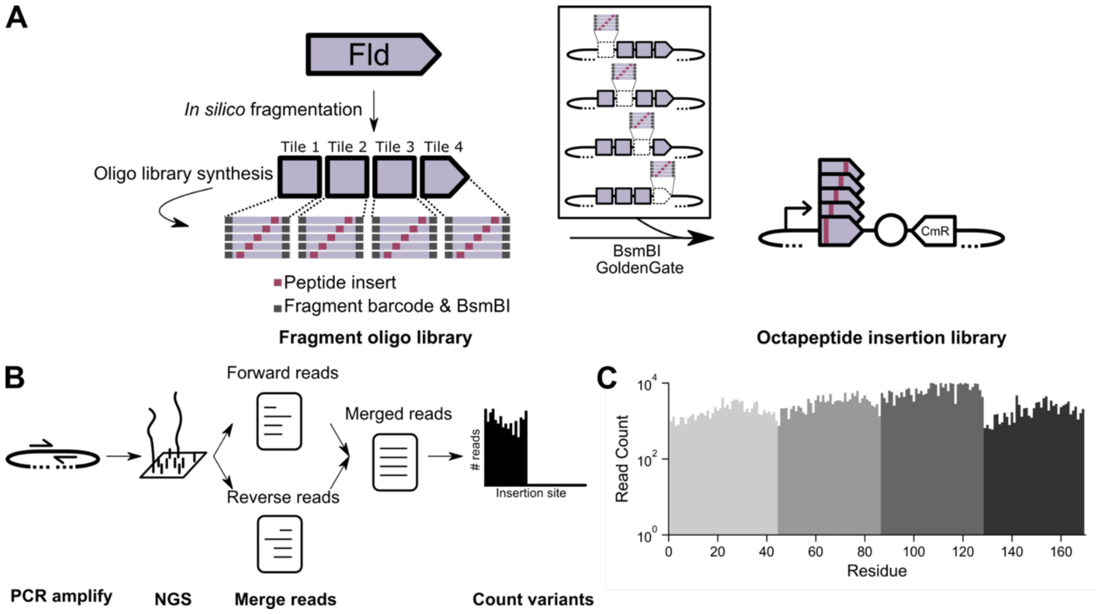


**Figure S3.** Fld library diversity following construction. (**A**) To create a vector library that contains every possible Fld variant arising from in-frame insertion of the oligonucleotide AGCGGGAGACCGGGGTCTCTGAGC that encodes the peptide SGRPGSLS, oligonucleotides were designed and synthesized as tiles. After using PCR to amplify each tile ensemble, the products were cloned into a Fld expression vector. The final library, which contains all possible insertion variants, is obtained by pooling equal quantities of the four tile libraries. (**B**) Next generation sequencing workflow for quantifying abundance of each Fld variant. The region targeted for sequencing yields forward and reverse sequencing reads, which were merged prior to quantifying the abundances of each insertion variant. (**C**) Abundance of each insertion variant after cloning of the Fld insertion library. Each vector library was sequenced and determined to contain roughly equal numbers of each expected insertion variant. Each unique color denotes unique tile library.

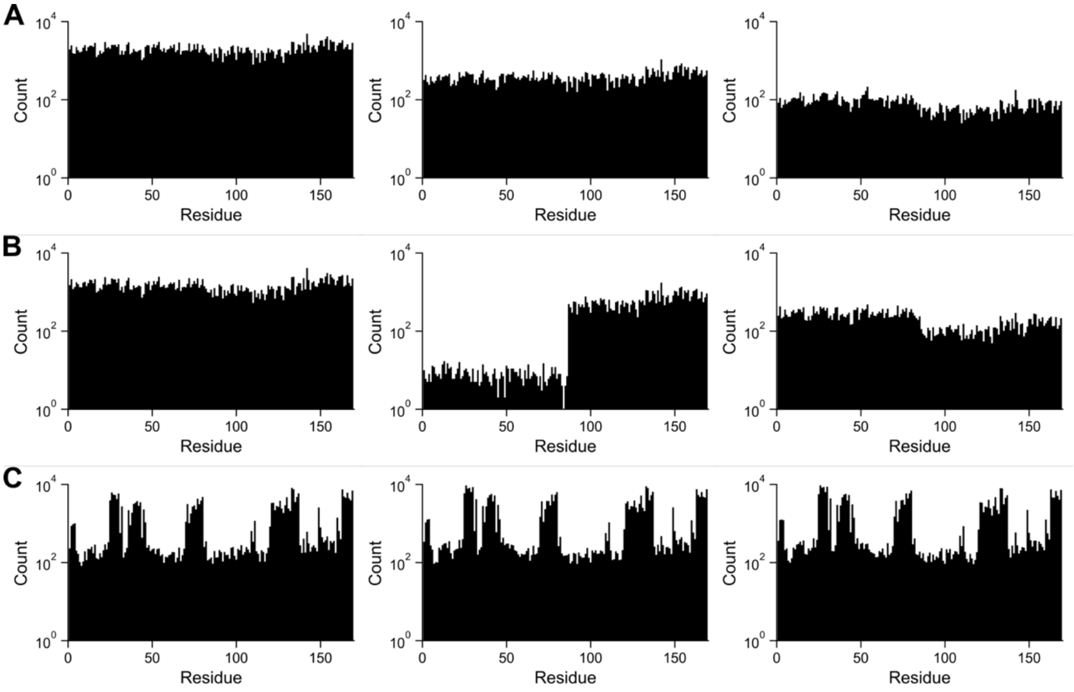


**Figure S4.** Fld insertion variant abundances for individual library samples in the (**A**) naive (**B**) transformed, and (**C**) selected libraries. The sequencing run which presented bimodal sequencing abundances was not used for calculating average abundances.

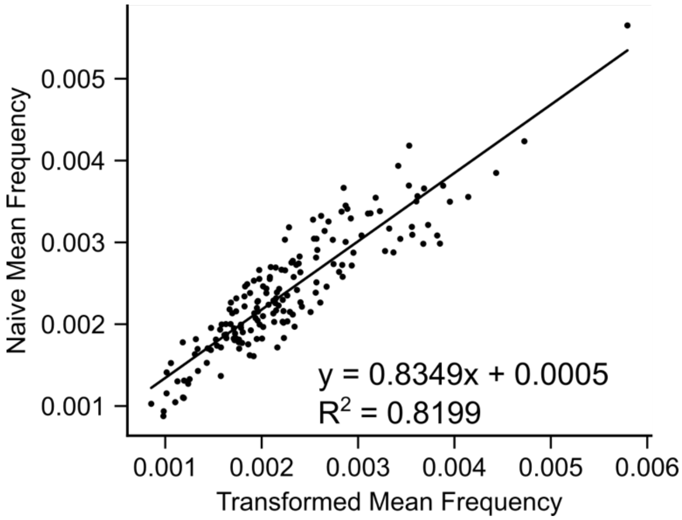


**Figure S5.** Relative frequencies of each variant in the naïve and transformed libraries. The mean frequency of each insertion mutant in both libraries was plotted and fit using linear regression.

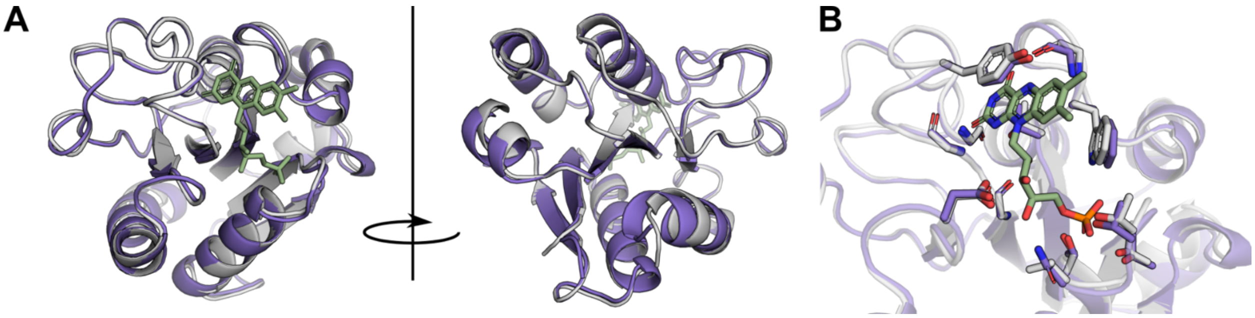


**Figure S6.** The structure of the *Synechocystis* Fld isiB predicted by AlphaFold (white, with FMN in green) from AlphaFoldDB was aligned with the crystal structure of *Synechococcus elongatus* Fld (PDB ID 1czl). Alpha carbon differ by 0.268Å RMSD.

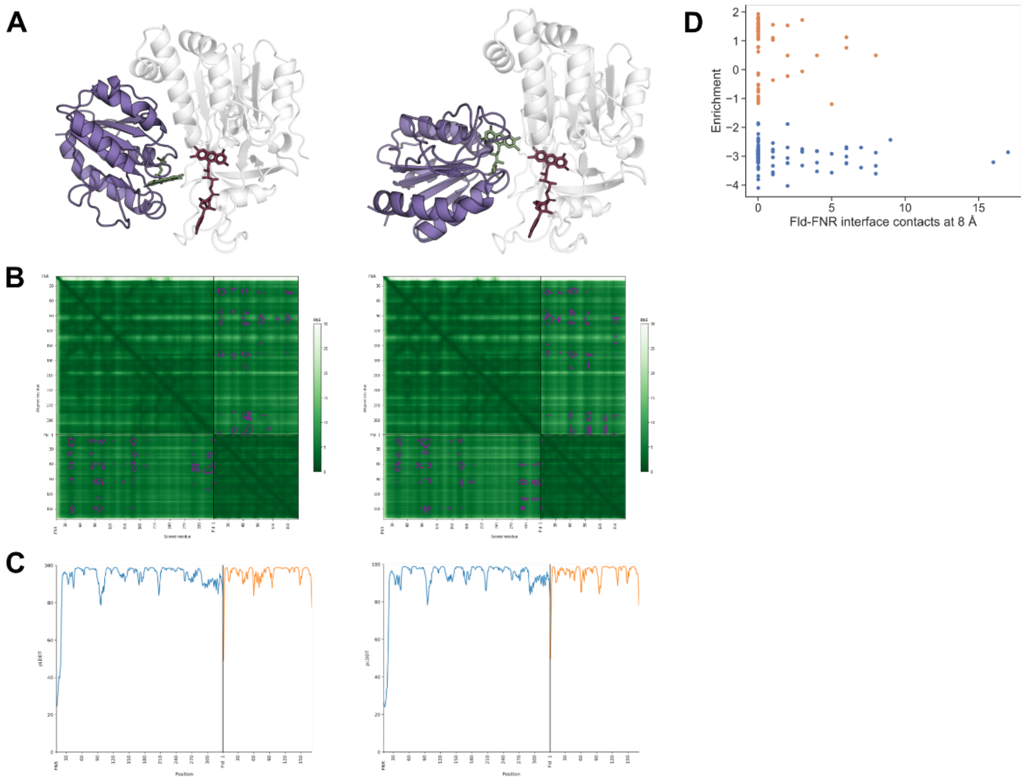


**Figure S7.** Structural modeling of the Fld-FNR complex. **(A)** AlphaFold predicts two binding modes for the Fld-FNR complex. Cofactors were placed by backbone alignment with the crystal structures 1czl and 1jb9. **(B)** Predicted aligned errors for the predicted binding modes. The vlues for pairs of residues containing atoms within 14 Å are outlined in magenta. **(C)** pLDDT plots for the Fld-FNR complexes. **(D)** Enrichments of Fld insertion variants plotted against numbers of intermolecular residue-residue contacts with FNR at 8 Å for the alternate binding mode.

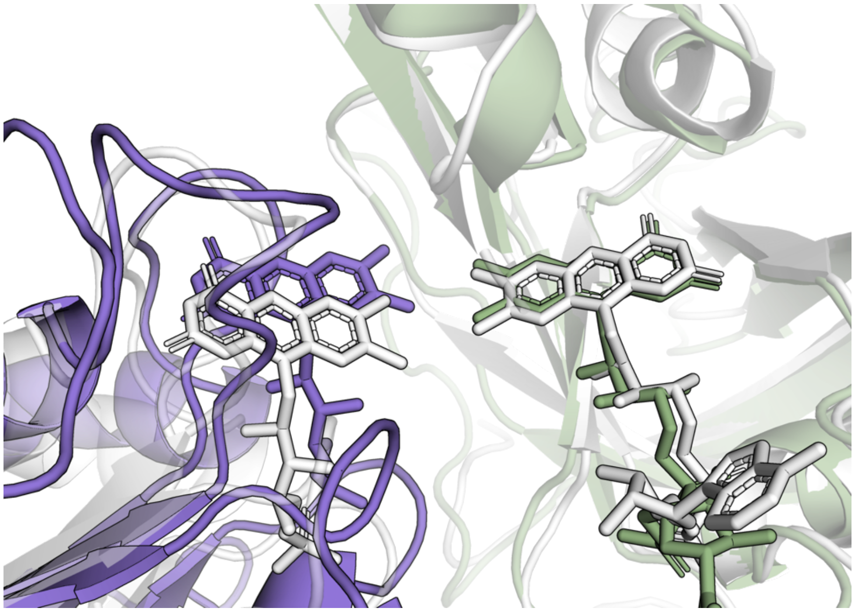

**Figure S8.** One of the predicted Fld-FNR binding modes closely matches the closed conformation of the Fld and FNR domains of cytochrome reductases (PDB ID 1amo, in white). Placement of FMN and FAD cofactors on Fld and FNR by backbone alignment with crystal structures (PDB IDs 1czl and 1jb9) reveals that the conformation predicted by AlphaFold recapitulates the 'side-to-side' geometry that supports interflavin electron transfer in cytochrome reductases.

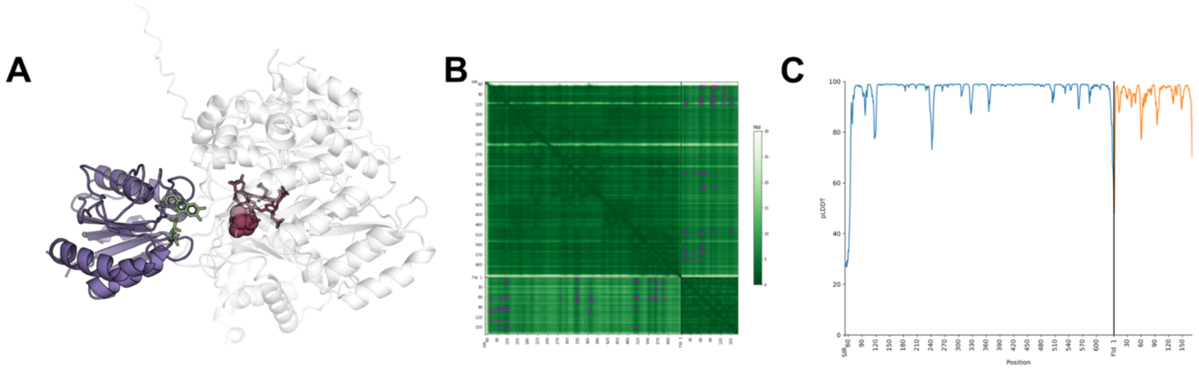


**Figure S9.** Structural modeling of the Fld-SIR complex. **(A)** The structure of the Fld-SIR complex predicted by AlphaFold and colored by enrichment. Cofactors were placed by backbone alignment with the crystal structures 1czl and 5h92. **(B)** Predicted aligned errors for the predicted complex. The values for pairs of residues containing atoms within 14 Å are outlined in magenta. **(C)** pLDDT plots for the Fld-SIR complex.

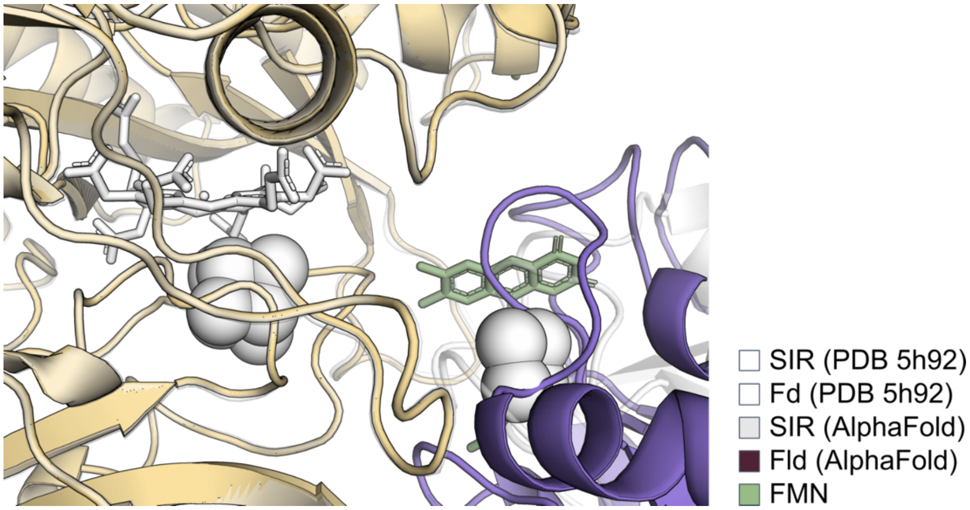


**Figure S10.** The inter-cofactor distance for the predicted Fld-SIR complex is <9 Å, which is conducive to intermolecular electron transfer. This is comparable to the intercofactor distances observed in crystal structures of Fd-SIR complexes.

**Table S1. Fld insertion variants (IV) chosen for characterization.** For each variant, the table indicates the variant name, residue prior to insertion, enrichment score, residues flanking insertion, and plasmid.

| Name | Residue | Enrichment | Flanking sites | Plasmid |
|---|---|---|---|---|
| IV-4 | I4 | -0.12025 | MTKI-GLFYG | pAT002 |
| IV-13 | G13 | -2.87361 | GTQTG-NTETI | pAT003 |
| IV-16 | E16 | -2.81574 | TGNTE-TIAEL | pAT004 |
| IV-27 | G27 | 1.580631 | QKEMG-GDSVV | pAT005 |
| IV-35 | M35 | -3.2111 | VVDMM-DISQA | pAT006 |
| IV-41 | D41 | 1.724116 | ISQAD-VDDFR | pAT007 |
| IV-43 | D43 | 1.558468 | QADVD-DFRQY | pAT008 |
| IV-48 | Y48 | -2.50062 | DFRQY-SCLII | pAT009 |
| IV-102 | A102 | -2.8928 | NFQDA-MGILE | pAT010 |
| IV-123 | T123 | 1.119366 | GFWPT-AGYDF | pAT011 |
| IV-127 | D127 | 1.408197 | TAGYD-FDESK | pAT012 |
| IV-129 | D129 | 1.309156 | GYDFD-ESKAV | pAT013 |
| IV-137 | G137 | 1.546849 | AVKNG-KFVGL | pAT014 |
| IV-143 | A143 | -2.79615 | FVGLA-LDEDN | pAT015 |
| IV-145 | D145 | -2.7498 | GLALD-EDNQP | pAT016 |
| IV-156 | R156 | -2.90211 | LTELR-VKTWV | pAT017 |
| IV-168 | L168 | 1.440814 | IKPIL-QS* | pAT018 |

**Table S2. Plasmids used for growth complementation measurements.** For each plasmid, the table notes the selection marker, origin type, and protein expressed. All of these vectors use aTc-inducible systems to regulate protein expression, with the exception of pSAC01, which uses constitutive promoters. IV, insertion variant.

| Name | Description |
|---|---|
| pAG034 | CmR, ColE1, Fld from *Synechocystis* PCC 6803 |
| pAG036 | CmR, ColE1, Fld from *Nostoc* PCC 7120 |
| pFd007 | CmR, ColE1, Fd from *Mastigocladus laminosus* |
| pFd007-C42A | CmR, ColE1, C42A mutant of *Mastigocladus laminosus* Fd |
| pSAC01 | SmR, p15A, FNR and SIR from *Zea mays* |
| pAT001 | CmR, ColE1, *Synechocystis* Fld with synonymous mutations |
| pAT002 | CmR, ColE1, *Synechocystis* Fld IV-4 |
| pAT003 | CmR, ColE1, *Synechocystis* Fld IV-13 |
| pAT004 | CmR, ColE1, *Synechocystis* Fld IV-16 |
| pAT005 | CmR, ColE1, *Synechocystis* Fld IV-27 |
| pAT006 | CmR, ColE1, *Synechocystis* Fld IV-35 |
| pAT007 | CmR, ColE1, *Synechocystis* Fld IV-41 |
| pAT008 | CmR, ColE1, *Synechocystis* Fld IV-43 |
| pAT009 | CmR, ColE1, *Synechocystis* Fld IV-48 |
| pAT010 | CmR, ColE1, *Synechocystis* Fld IV-102 |
| pAT011 | CmR, ColE1, *Synechocystis* Fld IV-123 |
| pAT012 | CmR, ColE1, *Synechocystis* Fld IV-127 |
| pAT013 | CmR, ColE1, *Synechocystis* Fld IV-129 |
| pAT014 | CmR, ColE1, *Synechocystis* Fld IV-137 |
| pAT015 | CmR, ColE1, *Synechocystis* Fld IV-143 |
| pAT016 | CmR, ColE1, *Synechocystis* Fld IV-145 |
| pAT017 | CmR, ColE1, *Synechocystis* Fld IV-156 |
| pAT018 | CmR, ColE1, *Synechocystis* Fld IV-168 |

**Table S3. Sequencing adaptors.** Sequences of the adaptors added to libraries prior to deep sequencing, which are amended to the ends of Fld gene amplicons.

| Direction | Sequence |
|---|---|
| Forward | 5'-ACACTCTTTCCCTACACGACGCTCTTCCGATCT-3' |
| Reverse | 5'-GACTGGAGTTCAGACGTGTGCTCTTCCGATCT-3' |

**Table S4. AlphaFold confidence metrics.** pTM, ipTM, Fld pLDDT, and Partner pLDDT scores for Fld complex models, including the two binding modes of the Fld-FNR complex (1) and (2).

| Complex | pTM | ipTM | Fld pLDDT | Partner pLDDT |
|---|---|---|---|---|
| *Fld-FNR (1)* | 0.91 | 0.86 | 93.5 | 95.8 |
| *Fld-FNR (2)* | 0.89 | 0.82 | 93.4 | 95.6 |
| *Fld-SIR* | 0.93 | 0.86 | 94.5 | 95.9 |
